# *Ex Vivo* Expanded Regulatory T Cells Inhibit AAA Progression by Limiting CD4+ and CD8+ T Cell Accumulation in Aortic Tissue

**DOI:** 10.1101/2025.10.08.681290

**Authors:** Chandrashekhar Dasari, Jose Luis Lopez, Masood Shawyan Jan, Sonali Shaligram, Pei-Yu Lin, Dina Lévy-Lambert, April Gu Huang, Carlos Lizama Valenzuela, Pierce Hadley, Qizhi Tang, Adam Oskowitz

## Abstract

**Background:** Regulatory T cells (Tregs) play a crucial role in the pathophysiology of abdominal aortic aneurysms (AAA), a chronic inflammatory condition with few treatment options for patients with early-stage disease. Treg therapy for AAA is potentially beneficial but its specific mechanism requires further investigation for clinical applications.

**Methods:** After identifying the critical role of T-cells in AAA using human and mouse AAA single-cell RNA sequencing data, we investigated the influence of Tregs on immune cell infiltration within mouse AAA—specifically CD3+ T cells—using congenic transfer of Thy1.1 allelic donor mice Tregs into AAA-induced wild-type C57BL/6J mice. AAA progression was quantified with ultrasound and image micrometry. Tissues obtained on postoperative days 7–42 were analyzed with flow cytometry, qRT-PCR, Verhoeff-van Gieson staining, hematoxylin-eosin staining, and immunohistochemistry.

**Results:** CD3+ T cell population was profoundly elevated in the elastase induced AAA mouse model which was further used in the study. The AAA mice that received Treg cell therapy had less elastin degradation and aortic wall enlargement than their control counterparts. Donor Tregs were detected in draining lymph nodes even after five weeks, with characteristic expression of FOXP3 and CD25. Although donor Tregs were not detected in the aortic microenvironment, the pro-inflammatory cell population including CD4 and CD8 cells was reduced, compared to control mice.

**Conclusion:** Elevated T cell population aggravates inflammation and promotes AAA progression. Treg therapy impedes the recruitment of T cells into AAA tissue by colonizing the draining lymph nodes, thereby mitigating AAA progression. This study deepens our understanding of Treg stability, function, and potential as a promising therapy for early-stage aneurysms.

## 1. INTRODUCTION

Abdominal aortic aneurysms (AAAs) are characterized by persistent vascular degeneration and aortic wall remodeling. The well-described interplay between the immune system and AAA pathogenesis consists of a cascade of inflammation and immune cell infiltration accompanied by increased cytokine production, extracellular matrix degradation and vascular smooth muscle cell dysfunction.^1,2^ Surgical intervention remains the primary definitive treatment for patients with rapidly progressing or large AAAs, who are at significant risk of rupture, a usually fatal event. However, surgery presents significant risk, including 1-5% perioperative mortality, and a 19-24% risk of re-intervention if performed endovascularly.^3^ There are few non-surgical therapies for patients with ongoing AAA progression, and no approved therapies for patients with small AAAs.^1^

In AAA, inflammatory cells, including macrophages, neutrophils, T lymphocytes, B lymphocytes, and endothelial cells, infiltrate the aortic wall.^4^ CD3+ T cells constitute half of the infiltrating immune cell population within AAAs, fostering the inflammatory milieu.^4,5^ Regulatory T cells (Tregs), a specialized subset of T lymphocytes, maintain immune homeostasis by modulating the pro- and anti-inflammatory microenvironments.^6^ Tregs promote the balance between inflammation and self-tolerance by suppressing the proliferation of effector T cells and antigen presenting macrophages.^7^ Patients with AAA have either or both impaired function and lower numbers of Tregs.^8^

Previous studies using AAA mice models have highlighted the pivotal role of Tregs in orchestrating the immune response in AAA.^9–11^ A recent study indicates that adoptive transfer of Tregs inhibits angiotensin II–induced AAA in apolipoprotein E–deficient (ApoE^−/−)^ mice susceptible to atherosclerosis.^12^ Another study found that Treg cells significantly reduced the incidence of AAA formation likely by inhibiting the production of prostaglandin E2, which plays a crucial role in inducing localized inflammation.^9^ Treating angiotensin II-infused mice with IL-2/anti-IL-2 monoclonal antibody complex expanded endogenous Tregs, decreased aortic wall inflammation and protected against AAA formation.^13^ In addition, the depletion of the Treg population led to increased mortality from AAA in mice.^10^ Notably, a clinical study reported a decrease in Treg cell numbers and reduced FOXP3 expression levels in CD4+CD25+ cells in peripheral blood from patients with AAA, implying that impaired immunoregulation by Tregs may be involved in the pathogenesis of human AAA.^8^ Collectively, these findings suggest that boosting of endogenous regulatory immune responses could be a possible therapeutic approach to prevent AAA.

Although these studies demonstrate the critical role of Tregs in protecting against AAA, the role of Tregs in modulating other immune cells, especially T cells which contribute to the pathogenicity of the AAA, has not been rigorously evaluated. A deeper understanding of the immunological heterogeneity of AAA is vital to unlocking novel Treg-mediated therapeutic strategies. Therefore, we sought to further define the protective role of Tregs in AAA, which we hypothesized is centered on the regulation of CD3+ T cell infiltration. In this study, we compared the T-cell distribution of human AAA samples and different AAA mouse models with respective controls using publicly available single-cell datasets. We then tested the effect of exogenous Tregs on immune cells in AAA. Our findings provide the first experimental evidence that Tregs reduce AAA progression by inhibiting T cell accumulation within the AAA microenvironment without directly colonizing aortic tissue. These findings provide potentially valuable insights for the development of Treg-mediated AAA therapy.

## 2. MATERIALS AND METHODS

### 2.1. Retrieval and processing of public data

Data from publicly accessible single-cell RNA sequencing of the human aorta (GSE166676)^14^ and three frequently used mouse models, CaCl_2_ (GSE164678),^15^ elastase (GSE152583),^16^ and angiotensin (GSE221789)^17^ were extracted from the GEO database. The R package Seurat v4.1.0 was used for cell filtration, normalization, principal component analysis (PCA), variable gene identification, clustering analysis, and uniform manifold approximation and projection (UMAP) dimensional reduction. Low quality cells were filtered out using the dataset authors’ established methods. For the human data, cells with <200 or >2500 genes expressed and those with >25% mitochondrial content were filtered out. For the mouse data, cells with <200 or >4000 genes and those with >10% mitochondrial content were filtered out. The resulting data was log-normalized and PCA was performed for dimensionality reduction. Leiden graph-based clustering in PCA space identified cell populations, which were further visualized using UMAP and various Seurat functions to explore gene expressions. The Seurat function FindAllMarkers was used to identify markers for each cluster of cells, which were matched with putative cell-type-specific markers to map each respective cluster to a corresponding cell type.

### 2.2. Animal studies

Mouse studies were conducted at the University of California, San Francisco (UCSF), USA. Mice were housed in a pathogen-free facility. Male C57BL/6J (B6J, Strain #000664) and B6 PL-*Thy1*^*a*^/CyJ (B6 Thy1.1, Strain #000406) mice were purchased from The Jackson Laboratory at 6-8 weeks of age. The mice were fed autoclaved standard chow (MI Laboratory Rodent Diet 5001; Harlan Teklad). All animal procedures were performed according to protocols approved by the Institutional Animal Care and Use Committee (IACUC) of UCSF. All experiments were performed on 6-to 8-week-old male mice. The experiments were conducted according to NIH guidelines.

### 2.3. AAA Elastase model

AAA was induced by topical elastase application in 6–8-week-old mice as previously described ^12^. Two days before surgery, the mice were fed 0.2% BAPN, an irreversible lysyl oxidase inhibitor β-aminopropionitrile, drinking water (BAPN, Sigma#A3137-25g, USA). After anesthesia, followed by a 1-2 cm abdominal incision, the infrarenal abdominal aorta was isolated from the renal vein to the iliac bifurcation. The infrarenal aorta was exposed to 5μL of either 100% porcine pancreatic elastase (#E1250, Sigma-Aldrich, St. Louis, MO, USA) or heat-inactivated elastase (100 °C, 3 min) for 5 minutes. After elastase exposure, the incision was stitched with PDS-II 5.0 absorbable sutures. Aneurysm progression was evaluated using weekly ultrasound imaging.

### 2.4. Cell culture

CD4+ cells from mouse axillary and inguinal lymph nodes (LNs) were extracted and separated using a negative selection kit (Stem Cell Technologies, Cat no#17952). CD4^+^CD25^**hi**^CD62L^+^ cells were purified by fluorescence-activated cell sorting (FACS) using a BD FACSAria Fusion Flow Cytometer. Tregs were cultured in DMEM supplemented with 10% FBS, 3000U/ml U/mL IL-2, and 1μM BME. Cells were stimulated with AntiCD3/anti CD28 coated dyna beads (Gibco#11452D) at a 1:3 ratio for 2 days and reactivated after 5 days. Mouse FOXP3 levels were checked every five days to maintain >90% purity of FOXP3^+^ cells.

### 2.5. Flow cytometry

For the sham group we combined three mice abdominal aortas for each sample and at least three samples were used for flow cytometry for each group. The aortas were digested using aorta digestion buffer to prepare a single cell suspension.^18^ The cells were resuspended in a flow buffer (DPBS with 1% bovine serum albumin) and incubated with surface antibodies for 30 min at 4 °C. For intracellular staining, cells were fixed and permeabilized with the eBioscience™ FOXP3 Staining Buffer Set (Cat#00-5523-00), as recommended by the supplier. The mouse FOXP3 antibody was added to the cells in the permeabilization buffer and incubated at room temperature for 30 min. The cells were washed with flow buffer and analyzed using a BD FACSAria Fusion Flow Cytometer.

### 2.6. Quantitative reverse transcription PCR (qRT-PCR)

Total RNA was extracted from snap-frozen aorta samples using RNA-later reagent following the manufacturer’s instructions (Sigma-Aldrich#R0901). cDNA synthesis was performed using the QuantiTect Reverse Transcription Kit (Qiagen#205313). Subsequent qPCR was done on an ABI Studio 7500 (Applied Biosystems) using the TaqMan™ Fast Advanced Master Mix (Taqman#4444556). Mouse *Gapdh* and Hprt were used as a reference genes to normalize sample amplification.

### 2.7. Hematoxylin and eosin (H&E) and elastin staining

The harvested tissues were snap-frozen using OCT (Leica, #14020108926) and sectioned using a cryotome at a thickness of 10 μm. On the day of staining, the sections were air-dried for 30 min, followed by H&E staining (VWR, #517-28-2) and Verhoeff-van Gieson staining (VVG) (Sigma-Aldrich, #1159740002) according to the manufacturer’s instructions. For each mouse, at least two sections were imaged and analyzed using a Nikon 6D high throughput color imaging microscope at magnifications of 10x and 20x. An unbiased and blinded manual grading system was used to assess the elastin degradation.^19^

### 2.8. Immunofluorescence

OCT tissue sections were air-dried for 15 min, followed by three 5-minute washes with phosphate-buffered saline pH7.4 (PBS). Subsequently, the sections were fixed in a 1:1 ratio of - 20 °C chilled methanol to acetone for 10 min. After three PBS washes, the tissues were blocked with blocking buffer (Cat# SC516214, Santa Cruz, CA, USA) for 1 h. A mixture of CD206 (Cat # AF2535, Novus biologicals, USA) and CD68 (Cat# 14-0688-82, Invitrogen, USA) antibodies were incubated overnight at a 1:200 ratio. After two PBS washes, the secondary antibodies were added and incubated for 2 h at room temperature in the dark. The tissues were dried briefly and fixed in DAPI-mixed anti-fade mounting medium (DAPI-Vecta mount, Cat# H-1200-10). At least two sections for each mouse were imaged at either 10x or 20x for analysis. Images were analyzed with FujiJ software.

### 2.9. Statistical analysis

Data entry and analysis were performed using Microsoft Excel 2010 and GraphPad Prism 8.0. Continuous variables are expressed as means and standard deviations (mean ± SD). Categorical data are presented as frequencies and percentages using the chi-square test. Differences between values were analyzed using nonparametric Mann-Whitney or Kruskal-Wallis tests and were considered significant at *p* <0.05.

## 3. RESULTS

### 3.1. Comparison of AAA immune cell heterogeneity of human vs mouse models

To identify critical immune cell populations within human aortic tissue, human immune cell clusters were generated using scRNA data from aneurysmal aortic tissue samples.^20^ We performed a Wilcoxon rank-sum test based on normalized data to identify marker genes in each cluster. Unsupervised Seurat-based clustering identified eight distinct populations (Figure 1A). Identities were assigned to each population based on gene expression patterns of established canonical markers of SMCs (SMC: *Myh11, Acta2)*, endothelial cells (EC: *Cdh5, Pecam1)*, fibroblasts (Fibro: *Col*1*a*1), macrophages (*Lyz2)*, neutrophils (*S100a8* and *S100a)*, dendritic cells (*Klrd1* and Flt3), T cell (*Cd3g)*, and natural killer cells (*Gzma)* and B-cells (*Cd79a, Ms4a1)*.^21,22^

**Figure 1:**
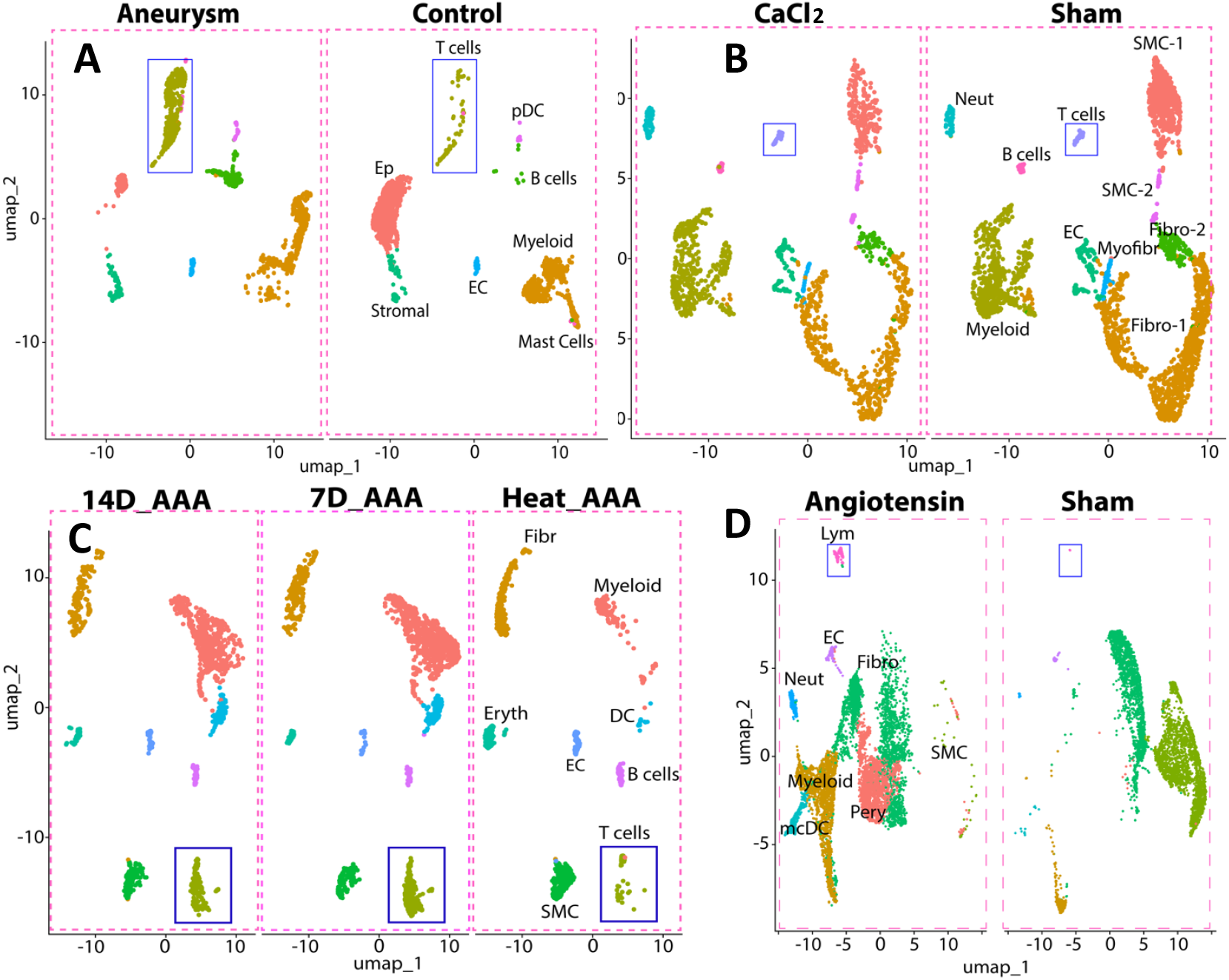
Immune cell clustering of the human and mouse AAA single-cell RNA sequencing data. A. Human AAA compared with healthy aortic tissues, B. Calcium chloride (CaCl_2_) treatment-induced AAA mouse aortic samples, C. Elastase-induced AAA mouse aortic samples, D. Angiotensin-II-infused AAA mouse aortic samples. Abbreviations: Sham= Saline or PBS injected, AAA=Abdominal aortic aneurysm, Heat_AAA=mouse aorta treated with heat inactivated Elastase, 7D_AAA=7days after AAA surgery, 14D_AAA=14days after AAA surgery. Cell clusters: T cells=CD45+ CD3+, Lym= Lymphocytes (CD45+); Fibr= Fibroblasts, Ep=epithelial cells; EC= Endothelial cells; pDC=peripheral dendritic cells; SMC= Smooth muscle cells; Neut= Neutrophils, Pery= Peripheral cells.

To identify a mouse model that exhibits a T-cell profile comparable to that found in human AAA tissue, we examined transcriptomic data and cellular composition profiles from three different mouse models (CaCl_2_, elastase, and angiotensin-II) retrieved from the GEO database and compared them to human aneurysmal aortic tissue. Interestingly, a 10-fold increase of T cells was observed in the elastase-induced mouse model (Figure 1C), whereas the CaCl_2_ model (Figure 1B) showed minimal or no change in the T cell population (Figure S1). The angiotensin-II mouse model did not exhibit a significant number of T cells during clustering (Figure 1D), even though a 10% mitochondrial cut-off was used, as opposed to other studies that used a 20% mitochondrial gene cut-off. This analysis indicates that the elastase-induced mouse model is the most promising candidate for further investigation of T-cell biology in relation to AAA disease because it closely resembles the T-cell profile found in human AAA tissue (Figure 1A).

### 3.2. Evaluation of T-cells during AAA formation and progression in the elastase-induced mouse model

The periadventitial elastase model of AAA is relatively new but has the benefit of recapitulating the chronic growth seen in human aneurysms.^23^ We knew of no study that had rigorously evaluated the long-term engraftment of T-cells in this mouse model and therefore first sought to define the increase in aneurysm-associated T-cells that we observed with single-cell data.

Heat-inactivated elastase was used as a negative control. Aortic enlargement became evident on postoperative day (POD) 7, as measured using B-mode ultrasound, with a significant difference in maximum aortic diameter compared to negative controls. Maximum aortic diameter continued to expand until day 42 (Figure 2A).

**Figure 2:**
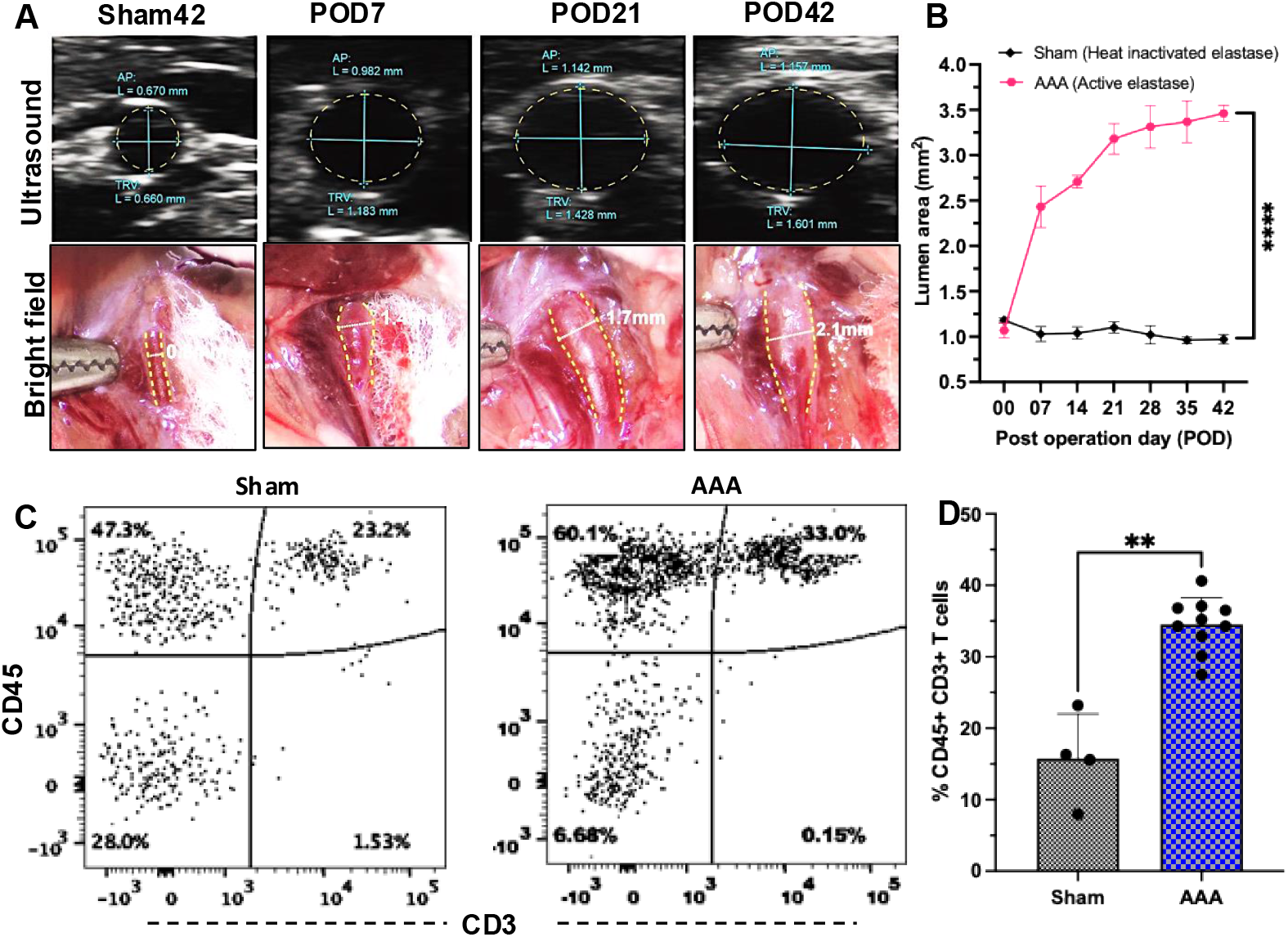
Evaluation of the elastase-induced AAA mouse model. A. Ultrasound imaging of the abdominal aorta and bright field microscopic images showing the lumen and outer aortic diameter on postoperative day7 (POD7), POD21, POD42; the POD42 of sham is represented as baseline control. B. Graphical representation of the lumen area calculated from ultrasound images. Lumen area = π(average diameter of AP + TRV)^2^; AP=anteroposterior and TRV = transverse dimensions. C&D. Representative flow cytometry images and graph of digested POD42 infra renal aorta showing CD45+ CD3+cells. Each group included at least 3 mice (n=3-9). Statistical analysis was performed using two-way ANOVA (Figure 2B) and the non-parametric Mann-Whitney test (Figure 2C). Significance depicted here as *p*<0.0001=****, **<005.

Images of the abdominal aortas were acquired during the harvesting process, and the external diameter was measured using image micrometry to corroborate ultrasound findings, as illustrated in Figure 2A. The results of the image micrometry were consistent with ultrasound measurements and showed substantial enlargement of the aorta in AAA mice compared to sham mice from POD7 to POD42.

To evaluate the correlation between the accumulation of CD3+ T cells and AAA, infrarenal abdominal aortas were digested in an enzymatic solution to prepare a single-cell suspension and analyzed for CD45+ CD3+ T cell populations using flow cytometry. The percentage of CD45+ CD3+ T cells (mean=41%) was significantly higher in the AAA group than in the sham (mean=14%) (p=0.001) (Figure 2B and 2D). Additionally, the overall cell population within the aortic tissue also shifted substantially, with the percentage of non-lymphocyte (CD45^-ve^) cells decreasing significantly from 28% to 6%, suggesting a preponderance of immune cells in the AAA samples.

### 3.3. Treg cell therapy reduces aortic enlargement

To facilitate cell tracking, Tregs to be used for adoptive transfer were isolated from the lymph nodes of Thy1.1 B6 mice using CD4+ negative selection and enriched for CD4^+^CD25^(high)^ CD62L^+^ cells using FACS and cultured *in vitro* (Figure 3A). On POD2 after AAA surgery or sham procedure, based on previous reports,^9^ 2 × 10^6^ Treg cells were administered via retro-orbital injection. The effect of Treg cell therapy on the aortic diameter was assessed via ultrasound every week starting on POD7 (5 days after cell injection) until harvest on POD42 (Figure 3B).

**Figure 3:**
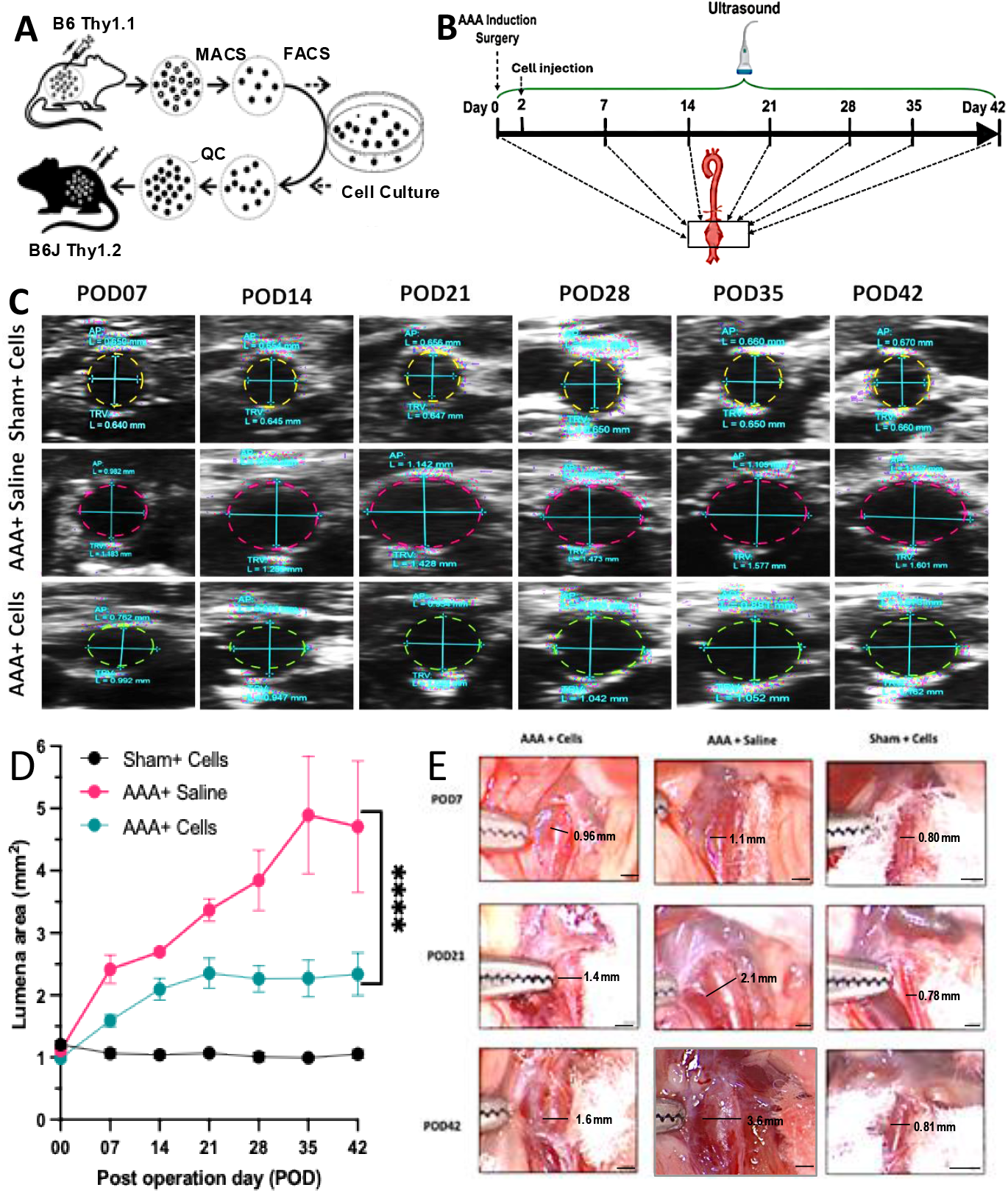
Treg cell therapy inhibits AAA progression. A&B) Schematic for adoptive cell transfer in a mouse model of AAA, C) Ultrasound showing the lumen of aorta, D) Mice undergoing AAA induction displayed up to 4.2mm^2^ of lumen area compared to 0.2mm^2^ of sham mice, Treg injected AAA mice displayed 2mm^2^, E) Outer diameter of the infrarenal aorta of sham, AAA, and Treg injected AAA mice measured using image micrometry. In the sham group, n≥3 mice were used at each time point. For AAA and AAA+ cells, n≥4 mice were used. For calculating statistical significance two-way ANOVA-Tukey’s multiple comparisons test, with single pooled variance was used and the *p* value represents 0.0001=****.

As depicted in Figure 3C and 3D, although modest AAA formation was evident, aortic enlargement was significantly lower in mice that received Treg cell therapy than in AAA mice. AAA formation was less apparent and ceased in mice that received Treg cell therapy (lumen area<2.4 mm^2^) after POD21, in contrast to AAA mice that continued to progress until POD42 (lumen area >2.4 mm^2^ to 5.8 mm^2^). Image micrometry showed the same trend: the external aortic diameters of Treg-injected AAA mice on POD21 and POD42 were smaller to those of their AAA counterparts (Figure 3E and Figure S2), suggesting the complete inhibition of AAA progression after POD21.

### 3.4. Treg cell therapy protects the aorta from inflammation damage

Damage to the extracellular matrix is a crucial factor in attracting pro-inflammatory immune cells. Structured elastin layering confers the elasticity to the aortic wall known to be damaged in AAA.^24,25^ VVG staining of the aortic tissues showed that elastin fiber layers in the abdominal aorta were significantly decreased and damaged in the AAA group compared to the sham group and the Treg injected AAA group (Figure S3A). Elastin degradation, assessed by an unbiased, blinded manual elastin degradation scoring system, showed that Treg treatment reduced elastin degradation (Figure S3B), which may be due to the protective role of Tregs in tissue damage.

H&E staining revealed the structural integrity of the abdominal aorta in the sham group, displaying a smooth outer membrane, wavy elastic fibers in the mid-membrane, and dense interlayer cells. In contrast, the AAA group exhibited degradation or rupture of the mid-membrane elastic and collagenous fibrous layers, along with significant infiltration of inflammatory cells into the abdominal aorta (Figure S3C). Furthermore, aortic wall thickness was significantly lower in the Treg-injected AAA group than that in the AAA group (Figure S3D).

Macrophages are tissue-infiltrating immune cells that produce matrix metalloproteinases such as MMP-1, MMP-2, and MMP-9, which degrade the extracellular matrix and damage aortic integrity, attracting further immune cell recruitment.^25^ Regulatory T cells reportedly suppress macrophages by inducing their differentiation into anti-inflammatory macrophages.^9^ In line with these previous reports our findings from immunostaining and qRT PCR showed that Treg cell administration resulted in macrophage polarization, shifting from pro-inflammatory M1 macrophages (identified by the CD68 marker) to anti-inflammatory M2 macrophages (identified by the CD206 marker), whereas AAA caused the accumulation of CD68-expressing macrophages (stained red in color) at higher levels (Figure S4).

### 3.5. Viability and detection of *in vitro-expanded* donor Tregs in recipient mice

To better understand the dynamics of donor Tregs, we conducted flow cytometry on the aortic tissue and draining lymph nodes of mice at POD42, with and without Treg cell therapy. We detected donor CD4+ CD8^-ve^ Thy1.1+ cells in the aortic and mesenteric lymph nodes of Thy1.2 recipient mice. However, the donor Thy1.1 Tregs were below the detectable levels in abdominal aorta. The percentage of Thy1.1 Tregs that survived varied among the mice, ranging from 0.04 to 0.2% of the mouse lymphatic T-cell population. Additionally, these cells consistently exhibited high levels of CD25, a characteristic marker of Tregs. The detected donor Tregs were CD4+ and not CD8+ (Figure 4).

**Figure 4:**
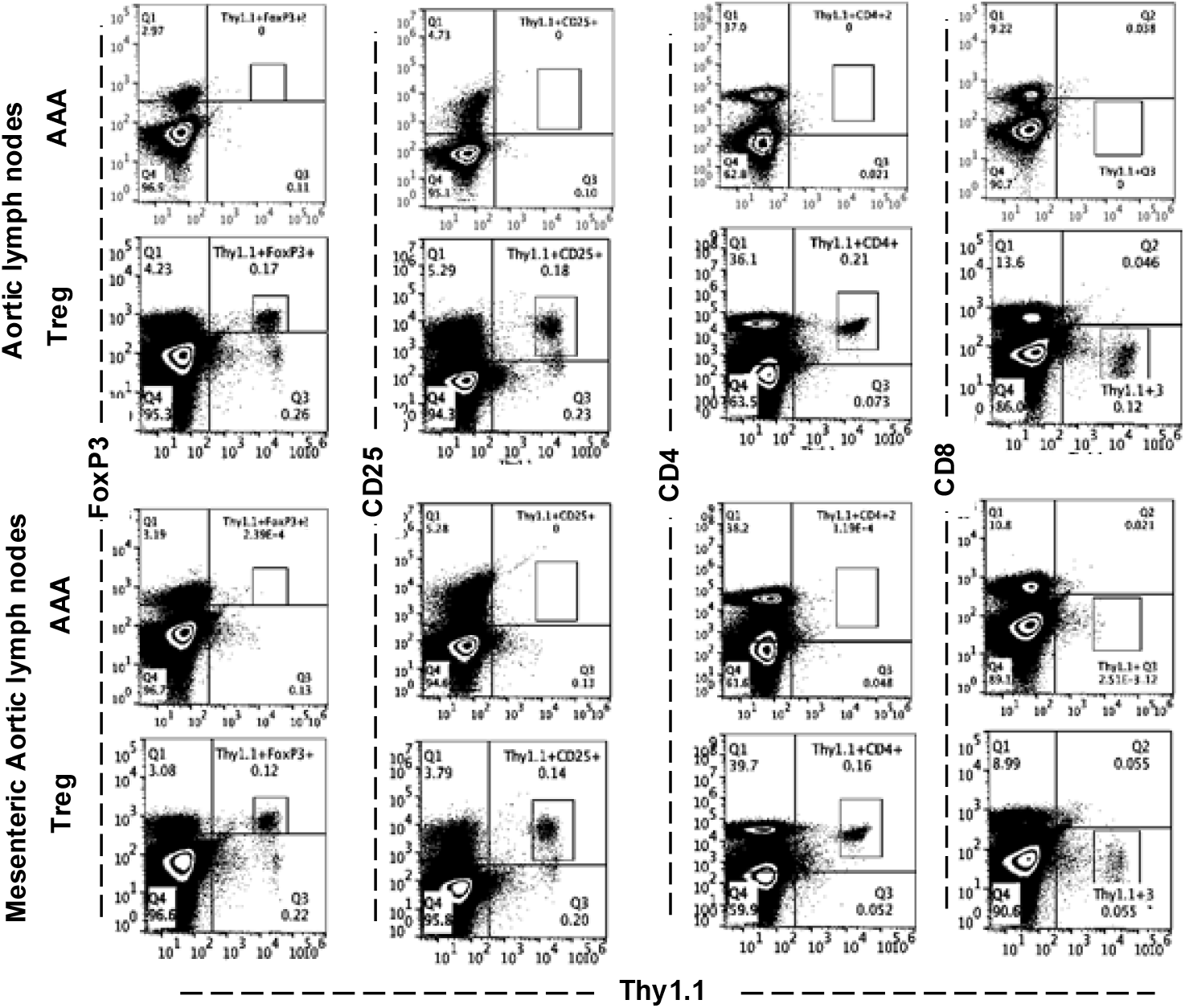
Analysis of donor Tregs (Thy1.1+ FOXP3+) in the draining lymph nodes. Representative images from flow analysis of lymphatic cells collected from the aortic and mesenteric lymph nodes stained for Thy1.1 against FOXP3, CD25, CD4, and CD8 respectively. Each group contains 3 to 8 mice (n=3-8).

### 3.6. Treg cell therapy decreased the T cell population in the aortic tissue

The results of real-time RT-PCR analysis of aortic mRNA (Figure 5A and 5B) indicated that the levels of *Cd*4 and *Cd8* transcripts were elevated in mice with AAA group compared to sham and Treg-injected AAA groups. The expression levels of the *Cd*4 gene in AAA mice remained constant from POD7-42, whereas the levels of *Cd8* increased significantly on POD21 and decreased at POD42. In contrast, the Treg cell therapy mice showed an initial spike in *Cd*4 levels on POD7, which decreased in POD21 and POD42. Interestingly, the transcript levels of *Cd*8 were higher than those of *Cd*4 in AAA mice, which was significantly moderated by Treg cell therapy. This observation is in line with the CD3 T cell sub-clustering of human scRNA data that showed an increase in CD4 and CD8 levels, with a greater emphasis on CD8 (129 events) than CD4 (102 events) (Figure 5C).

**Figure 5:**
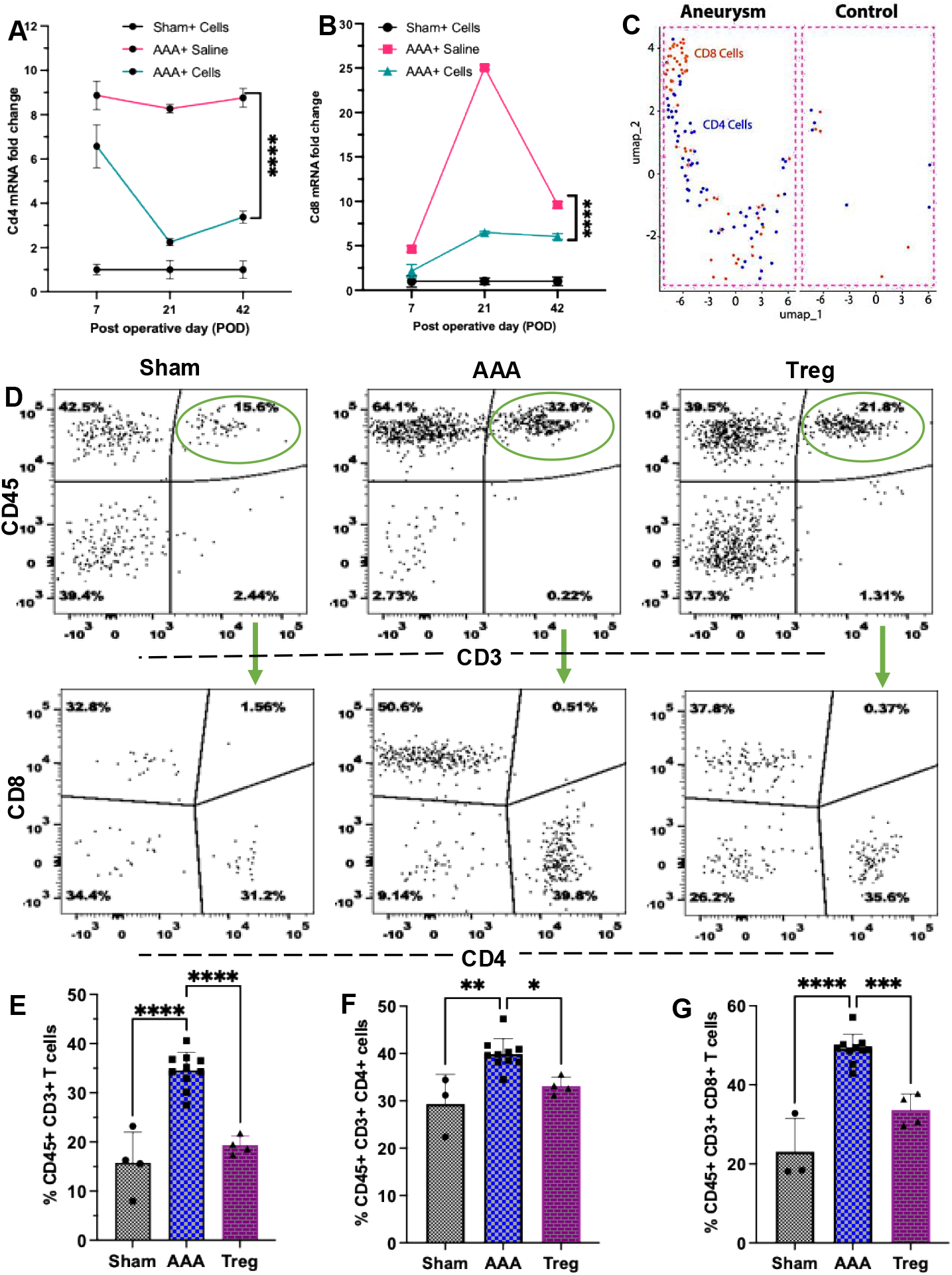
Treg cell therapy reduces tissue-resident T cells. A&B. qRT PCR results showing the *Cd*4 and *Cd*8*a* RNA levels in mouse abdominal aorta Sham, AAA and Treg groups, C. CD8 and CD4 cell clusters in human AAA samples and controls. D. Flow cytometric analysis of POD42 samples showing lowering of CD45+ CD3+ T cells by Treg cell therapy. E. Bar graph showing CD45+ CD3+ T cells in aortic tissues. F. Bar graph showing CD45+ CD3+ CD4+ T cells in aortic tissue. G. Bar graph showing CD45+ CD3+ CD8+ T cells in aortic tissues. Each group included at least 3 mice (n ≥ 3). Statistical analysis was done using two-way ANOVA (Figure A&B) and the non-parametric Mann-Whitney test (Figure 5C-G). *p* value depicted <0.0001=****, ***≤ 0.001, ** ≤ 0.005, * ≤ 0.05.

To validate the observations from qRT-PCR and in silico analyses, we analyzed resident T cells in aortic tissue in AAA mice with and without Treg cell therapy, as well as in sham mice. The T-cell population was exceptionally low in the abdominal aortic tissue of sham mice but accumulated significantly in AAA mice. In AAA mice, the population of CD45+CD3+ lymphocytes was double that of sham mice. In Treg-injected mice, the T-cell population increased modestly compared to the sham mice but were significantly less than the AAA mice. CD45+ CD3+ CD4+ cells were prevalent in AAA mice (∼42%) but not in Treg-injected AAA mice and sham group (both fall in the range of 30% to 36%). In addition, the accumulation of CD8+ cells in AAA condition was inhibited by Treg cell therapy and decreased to 37% compared to 55% in AAA mice.

## 4. DISCUSSION

The progression of AAA is strongly associated with chronic inflammation, which is exacerbated by the invasion of pro-inflammatory immune cells. Disruption of the immune system balance leads to tissue destruction and aneurysm formation.^26^ Although mouse models for studying AAA use Ang-II, elastase, or CaCl_2_ to chemically induce AAA, the rates of formation and rupture differ from those in human AAA.^27^ For instance, AAA induced using the Ang-II mouse model was more frequently complicated by intramural hematomas and aortic dissections than in human AAA.^28,29^ Using publicly available single-cell RNA data, we found that the T-cell makeup and rate of aneurysm growth in the elastase AAA mouse model was closest to that of human AAA.

Tregs, a specific subset of the adaptive immune system, are crucial for maintaining aortic wall stability.^8^ Rodent-based studies have suggested that Tregs can reduce or even prevent AAA development by neutralizing this damaging inflammation.^30^ Understanding Treg stability, expansion, and viability in the recipient body, particularly in association with other immune cells, is crucial for the successful transition of this approach into clinical use. To bridge this gap, we conducted experiments in recipient Thy1.2 allelic C57BL/6J mice with Thy1.1 allelic B6.PL- Thy1a/CyJ Treg donors. On postoperative day two, recipient mice that had undergone AAA induction using topical elastase administration were injected with in vitro expanded Tregs from donor mice. Ultrasound imaging calculating the luminal area showed that *ex-vivo* expanded Tregs significantly slowed AAA progression in mice in the first three weeks when compared to non-treated mice, and showed no further growth of AAA thereafter, whereas AAA progression continued in untreated mice. Significantly less elastin degradation was observed in Treg-treated AAA mice compared to controls. Histology showed slower aortic wall thickening in treated mice, with wall thickness plateauing after three weeks, in contrast to the continued growth seen in untreated mice through week six.

Activated M1 macrophages infiltrate the aortic tissue and secrete matrix-degrading enzymes (MMPs) that contribute to aortic-wall damage in AAA.^31,32^ Anti-inflammatory M2 macrophages are crucial to re-establish the immune balance after an inflammatory event.^31,33^ Treg cell therapy has been shown to improve the macrophage polarization from M1 to M2 and to co-infiltrate inflamed vascular tissue.^4,34^ We found increased polarization of the macrophage population from M1 to M2 in Treg-injected AAA, with the measured M1/M2 ratio being closer to that of control mice than of untreated AAA mice.

Our study also confirmed donor Treg viability and functionality in Thy1.2 recipient mice. CD4+ CD8^-ve^ Thy1.1+ expression was used as a signature for donor Tregs to identify them in the abdominal aorta-draining lymph nodes of recipient mice. Importantly, donor Tregs maintained elevated levels of CD25 expression at 6 weeks, which is characteristic of FoxP3+ CD4+ Tregs,^35^ demonstrating their survival and retention of full functionality in recipient mice.

CD3+ T lymphocytes, comprising CD4+, CD8+, γδ, and regulatory T cells, constitute >50% of lymphocytic infiltrate in AAA tissue,^4,36^ CD4+ T helper cells are predominant, with Th1 subsets secreting pro-inflammatory cytokines and Th2 subsets mediating anti-inflammatory responses.^36^ CD8+ tissue-resident T cells contribute to the pro-inflammatory milieu and are present in higher proportions relative to CD4+ T cells within AAA lesions.^36,37^ This T-cell distribution and functional heterogeneity significantly influences the inflammatory processes underlying AAA pathogenesis. Tregs suppress conventional T-cell proliferation by overexpressing IL2 receptor, anti-inflammatory cytokines, and inhibitory signaling molecules, which combats pro-inflammatory signals to re-establish immune homeostasis.^38^ Our study shows that Treg therapy suppressed the infiltration of CD3+ T cells (including both CD4+ and CD8+ subsets) in AAA wall and reduced the proportion of CD8+ T cells compared to CD4+ T cells. This was confirmed by flow cytometry and qRT-PCR. Collectively, our results demonstrate that Tregs modulate the accumulation of T cells and macrophages within the aortic wall, thereby serving a protective role against AAA.

## 5. LIMITATIONS AND CONCLUSIONS

An important limitation of our preclinical study is its reliance on a single rodent model. Using only one model cannot fully replicate the complex pathogenic environment of patients with AAA, including factors such as blood pressure, flow rate, and internal circulation composition. Further validation using larger animal models is essential before translating this Treg therapy into clinical practice.

In conclusion, our study shows that *ex vivo* expanded Tregs injected into AAA mice remain viable, functional, and inhibit AAA progression. These findings refine our understanding of the mechanism underlying Treg therapy for AAA and support its potential use for early stage or small AAAs, in which Treg therapy may prevent progression to more rupture-prone aneurysms.

Additionally, Treg therapy could be an option for patients with large aneurysms who are poor candidates for surgical intervention. Although *ex vivo* expanded Treg cell therapy is promising, key areas related to this type of research require further improvement, these include the loss of genetic stability due to *ex vivo* culture conditions, issues with freeze and thaw cycles, donor cell viability in the recipient, and overall costs. Finally, it is crucial to explore genetic engineering techniques to maximize the effectiveness of *ex vivo* expanded Tregs as immunotherapeutic agents in the clinical setting.

## 6. ASSOCIATED CONTENT

Supporting Information: Supplementary Figures S1-S5.

Table: List of amplification primers

Table2: Flowcytometry and immunohistochemistry antibody list

## 7. FUNDING STATEMENT & ACKNOWLEDGMENTS

This project was funded via grants from the NIH (1R21HL173856-01) and the AHA Innovative Project Award. This study was supported in part by HDFCCC Laboratory for Cell Analysis Shared Resource Facility through a grant from NIH (P30CA082103).

## 9. AUTHOR CONTRIBUTIONS

CD contributed to the implementation of the research, analysis of the results, and writing of the manuscript; JL, MJ, AH, SS, and PL contributed to the implementation of the research. PH and QT have helped to establish experimental protocols. AO conceived and supervised the project. All other authors helped in experimental planning, the analysis of the data and writing the manuscript.

## 11. ABBREVIATIONS

AAA: abdominal aortic aneurysm
Treg: regulatory T cells
Treg: CD3+ CD4+ CD62L+ CD25^**hi**^
ACT: advanced cell transfer
IACUC: Institutional Animal Care and Use Committee.

